# Thermodynamic versus kinetic basis for the high conformational stability of nanobodies for therapeutic applications

**DOI:** 10.1101/2024.08.28.609651

**Authors:** Atanasio Gómez-Mulas, Eduardo Salido, Angel L. Pey

## Abstract

Single domain nanobodies (NB) are powerful tools for biotechnological and therapeutic applications. They strongly bind to their targets and are very stable. Early studies showed that NB unfolding is reversible and can be analyzed by equilibrium thermodynamics whereas more recent studies focused on their kinetic stability in very harsh conditions, far from storage or physiological temperatures (4-37°C). Here we reinforce the thermodynamic view in which a simple two-state denaturation model is applicable. We found that thermal stability of NB actually reflect thermodynamic stabilities in wide range of temperatures (18-100°C). We also modeled their structure observing subtle differences. We expect that our approach will be helpful to improve our capacity to enhance structure-function-stability relationships of NB.

## Introduction

Protein stability is a key factor for therapeutic application of proteins since therapeutic proteins must retain their activity at *in vivo* conditions [1–3]. However, the relationships between *protein stability* (thermodynamic and kinetic stability, local unfolding…) and the shelf life of therapeutic proteins is not well understood. Here we address how the thermal stability of three nanobodies (NB) raised to correct protein misfolding in a rare genetic disease (primary hyperoxaluria type 1, [4–6]) correlate with thermodynamic stability in a wide range of temperatures.

NB are single-domain and antigen-specific fragments derived from the heavy-chain antibodies of camelids, such as alpacas, llamas and dromedaries, with a great potential as therapeutic agents [7–13]. NB bind to their targets through three complementarity-determining regions or CDR (CDR1, CDR2, and CDR3) that contain each a hypervariable loop or (HVL 1-3)[7]. These CDR (particularly CDR3) determine their high specificity and affinity for its target [7]. In an early study, it was described that NB unfolding is highly reversible at low temperatures and that NB display high conformational stability, with an unfolding free energy in the range of 10-15 kcal·mol^-1^ and resembling well a two-state equilibrium unfolding model [14]. Thus, considering simple Lumry-Eyring models, one could consider that the shelf life of NBs is associated with the thermodynamics (and kinetics?) of unfolding under storage (e.g. 4°C) or working (37°C) temperatures [15–17]. As expected, incubation of NB at very high temperatures [in the range 70-85°C, close to the apparent melting temperature (*T*_m_) of the NB] led to irreversible denaturation, and this approach was alternatively used to explain the different thermostability of different NB [18,19]. However, we must note that studies carried out at such a high temperatures imply rather long (kinetic) extrapolations to low temperatures, and that the irreversible processes operating at high temperatures may not hold at low temperatures [i.e. different irreversible processes may show different activation energies (*E*_a_) according to a simple Arrhenius analysis].

Protein stability studies derived from thermal denaturation experiments are not always easy to be carried out strictly and difficult to understand deeply [20]. Two key factors necessary to apply a suitable model to yield thermodynamic and/or kinetic properties from protein thermal denaturation scans are the experimental reversibility and the scan-rate dependence [20]. A high reversibility of the thermal transition(s) has been traditionally taken as a *bona-fide* proof for applying thermodynamic analysis to thermal denaturation experiments, although the scan-rate dependence must be also taken into account [20,21]. For instance, thermal denaturation of human phenylalanine hydroxylase has been shown to be irreversible but its scan-rate dependence vanishes at high scan rates (about 1°C·min^-1^) allowing partial characterization of its unfolding thermodynamics using equilibrium models as long as the post-transition behavior (highly distorted by irreversible processes) is not taken into account [22]. Importantly, early works carried out with a wide variety of NB showed that their reversibility upon thermal denaturation is moderate-high (typically 50-100%) when moderate scan-rates are used (0.5-1°C·min^-1^)[14,23,24], whereas other authors have described thermal denaturation of NB as a kinetically controlled process and not dictated by equilibrium thermodynamics [17,18]. Therefore, it is not clear how the high thermodynamic stability of NB at room (or storage or physiological) temperature is connected (or not) with a high thermal stability and which are the structural-energetic basis of their stability (thermodynamic and/or kinetic stability, local and global stability).

In this work, we have confirmed the relationships between equilibrium and thermal stability of different NB showing that their thermostability can be explained well from a higher thermodynamic stability at all temperatures, particularly at working conditions (around 37°C for a therapeutic NB). The binding affinity of these three NB (named NB-AGT-1, -2 and -6) vary in 3 orders of magnitude, *K*_d_ ranging from 3.8 to 5·10^−3^ nM to AGT-LM (the *minor* allele of alanine:glyoxylate aminotransferase associated with the rare disease name primary hyperoxaluria type I, OMIM # 259900; [25] [Gomez-Mulas, to be submitted]. We combined chemical and thermal denaturation experiments to generate thermodynamic stability curves for these NB that differ in 15°C in the thermal denaturation temperatures and about 10 kcal·mol-^1^ in the unfolding free energy at room temperature.

## Materials and methods

### Nanobody (NB) expression and purification

NB generation was carried out by NabGen Technologies (Marseille, France), and further details will be provided in a future publication [Gomez-Mulas, to be submitted]. *E. coli* BL21 (DE3) cells were transformed with the pET-24(+) vector containing the cDNA of each NB (NB-AGT-1, -2 and -6) and carrying a C-terminal 6His-tag. A preculture of 240 mL of LB medium containing 30 µg·mL^-1^ kanamycin (LBK) was inoculated with transformed cells and grown for 16 h at 37 ºC. These cultures were diluted into 4.8 L of LBK, grown at 37 ºC for 3 h and NB expression was induced by the addition of 0.5 mM IPTG (isopropyl β-D-1-thiogalactopyranoside) and lasted for 6 h at 25 ºC. Cells were harvested by centrifugation and frozen at -80 ºC for 16 h. Cells were resuspended in binding buffer, BB (20 mM Na-phosphate, 300 mM NaCl, 50 mM imidazole, pH 7.4) plus 1 mM PMSF (phenylmethylsulfonyl fluoride) and sonicated in an ice bath. These extracts were centrifuged (20000 g, 30 min, 4 °C) and the supernatants were loaded into IMAC (immobilized-metal affinity chromatography) columns (Cytiva, Barcelona, Spain), washed with 40 bed volumes of BB and eluted in BB containing a final imidazole concentration of 500 mM. These eluates were immediately buffer exchanged using PD-10 columns (Cytiva, Barcelona, Spain) to 50 mM HEPES [N-(2-Hydroxyethyl)piperazine-N’-(2-ethanesulfonic acid)]-KOH pH 7.4, analyzed by SDS-PAGE (Polyacrylamide gel electrophoresis in the presence of sodium dodecyl sulphate) and stored at -80°C upon flash freezing in N_2_. NB-AGT samples were further purified by loading them onto a SuperDex 75 16/60 size exclusion chromatography column (Cytiva, Barcelona, Spain) using 20 mM HEPES-NaOH, 200 mM NaCl pH 7.4 as mobile phase at 1 mL·min^-1^ flow rate. Fractions containing NB-AGTs were collected, concentrated, buffered exchange to 50 mM K-phosphate pH 7.4 and stored at -80 ºC after flash-freezing in liquid N_2_. Purity and molecular weight was analysed again by SDS-PAGE and further checked by high performance liquid chromatography coupled to electrospray ionization mass spectrometry (HPLC/ESI-MS) by the High-resolution mass spectrometry unit, Centro de Instrumentación Científica (University of Granada) in a WATERS LCT Premier XE instrument equipped with a time-of-flight (TOF) analyzer. NB-AGT concentration was measured using the following molar extinction coefficients (ε_280_) according to their primary sequence: NB-AGT-1.-25565; NB-AGT-2.-33015; NB-AGT-6.-20065, all in M^-1^·cm^-1^.

### Differential scanning calorimetry (DSC)

DSC experiments were carried out using a VP-DSC differential scanning microcalorimeter (Malvern Pananalytical, Malvern, UK) with a cell volume of 137 µL and automated sampling. Experiments were performed in 50 mM K-phosphate pH 7.4 using 20 µM of NB-AGT and typically at scan rate of 2°C·min^-1^ (even though other rates were tested to check for scan rate dependence). Scans were carried out in a temperature range of 20-90°C (NB-AGT-1 and NB-AGT-6) or 20-100°C (NB-AGT-2) to allow complete unfolding and to minimize distorsions from irreversible thermal denaturation. Actually, the reversibility is quite significant although not complete, thus supporting the applicability of equilibrium denaturation to determine some relevant unfolding parameters (reversibility was 47±1% for NB-AGT-1, 52±2 for NB-AGT-2 and 50±1% for NB-AGT-6; from calorimetric enthalpies of upscans and rescans shown in Figure 1). In some DSC experiments, a guanidium hydrochloride (GdmHCl) solution was prepared in 50 mM K-phosphate pH 7.4 by weight at ∼ 6 M and used at a final concentration of 0-0.9 M (all GdmHCl concentrations were checked by refractive index measurements).

**Figure 1.**
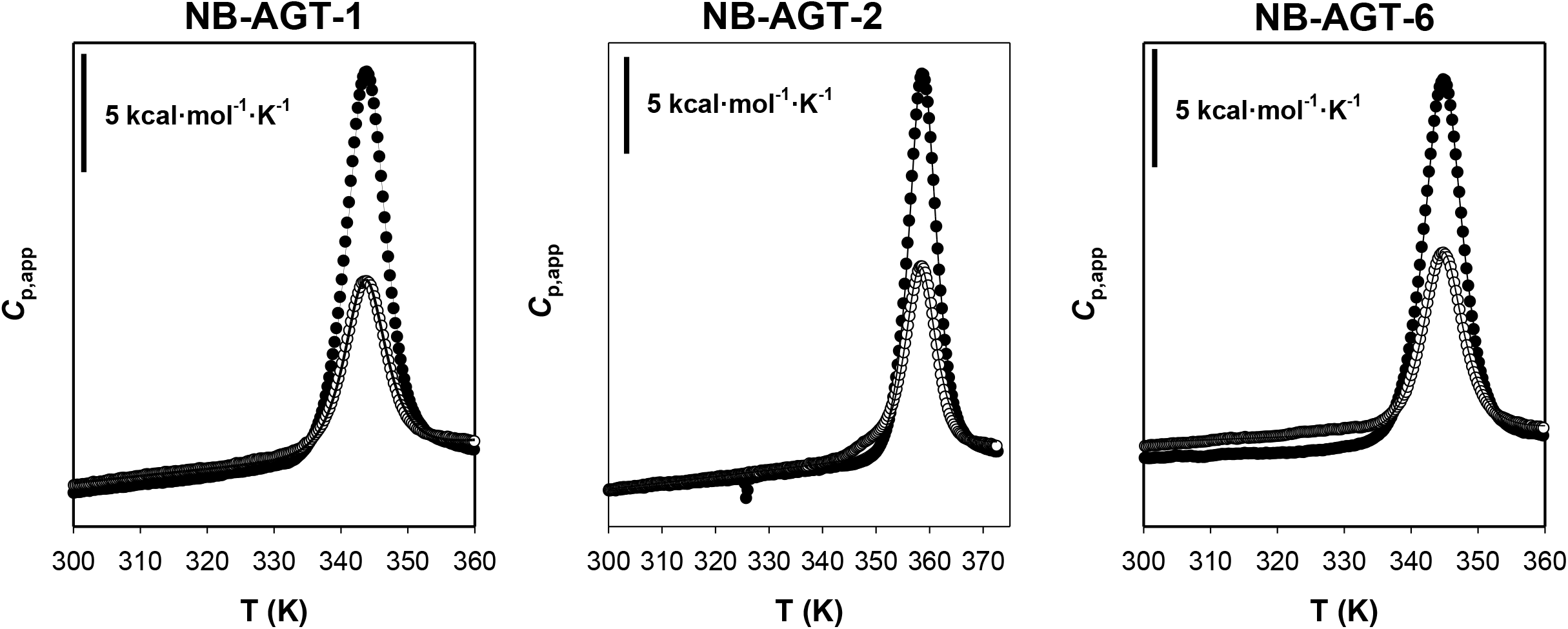
A two-state equilibrium thermal denaturation model provides a good description of the thermal unfolding of NB-AGT-1, NB-AGT-2 and NB-AGT-6. Closed circles show the upscans and the open circles the rescans. Lines are best-fit to equation 1. Data are from one experiment.

As a first approach to quantitatively evaluate the thermal denaturation behavior of NB-AGT, we applied a model to analyze the DSC scans in which equilibrium denaturation is not assumed to be a two-state process (i.e. not only the native and unfolded may be populated). To this end, we used a function in which the temperature (*T* in K) dependence of the apparent heat capacity (*C*_p,app_) was described by equation 1:

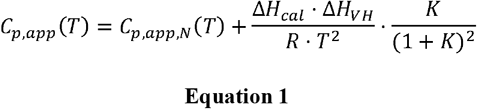

Where *C*_p,app,N_ is the temperature-dependent apparent heat capacity of the native state (described by a straight line equal to a+b·T), the area under the calorimetric *peak* or calorimetric enthalpy (Δ*H*_cal_), the Van’t Hoff enthalpy (Δ*H*_VH_) that describes the temperature-dependence of the denaturation equilibrium constant *K* (equation 2; the *T*_m_ is the temperature at which *K*=1) and *R* is the ideal gas constant (1.987 cal·mol^-1^·K^-1^).

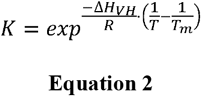

The unfolding free energy (Δ*G*) at a given temperature was calculated using the well known equation ΔG = -R·T·ln *K* at temperature within the thermal transition. We did not attempt to calculate the denaturation heat capacity (Δ*C*_p_) from DSC scans since the unfolded heat capacity might be significantly affected by the lack of complete reversibility (higher temperatures may largely enhance irreversible processes). Instead, we estimated an average Δ*C*_p_ for the three NB-AGT from the *T*_m_-dependent values of Δ*H*_cal_ in the absence or presence of GdmHCl (Figure 2). Please note that for fitting and display purposes, Temperature is shown as K or as °C in some figures (K is required for fittings but °C is more friendly for a broad readership).

**Figure 2.**
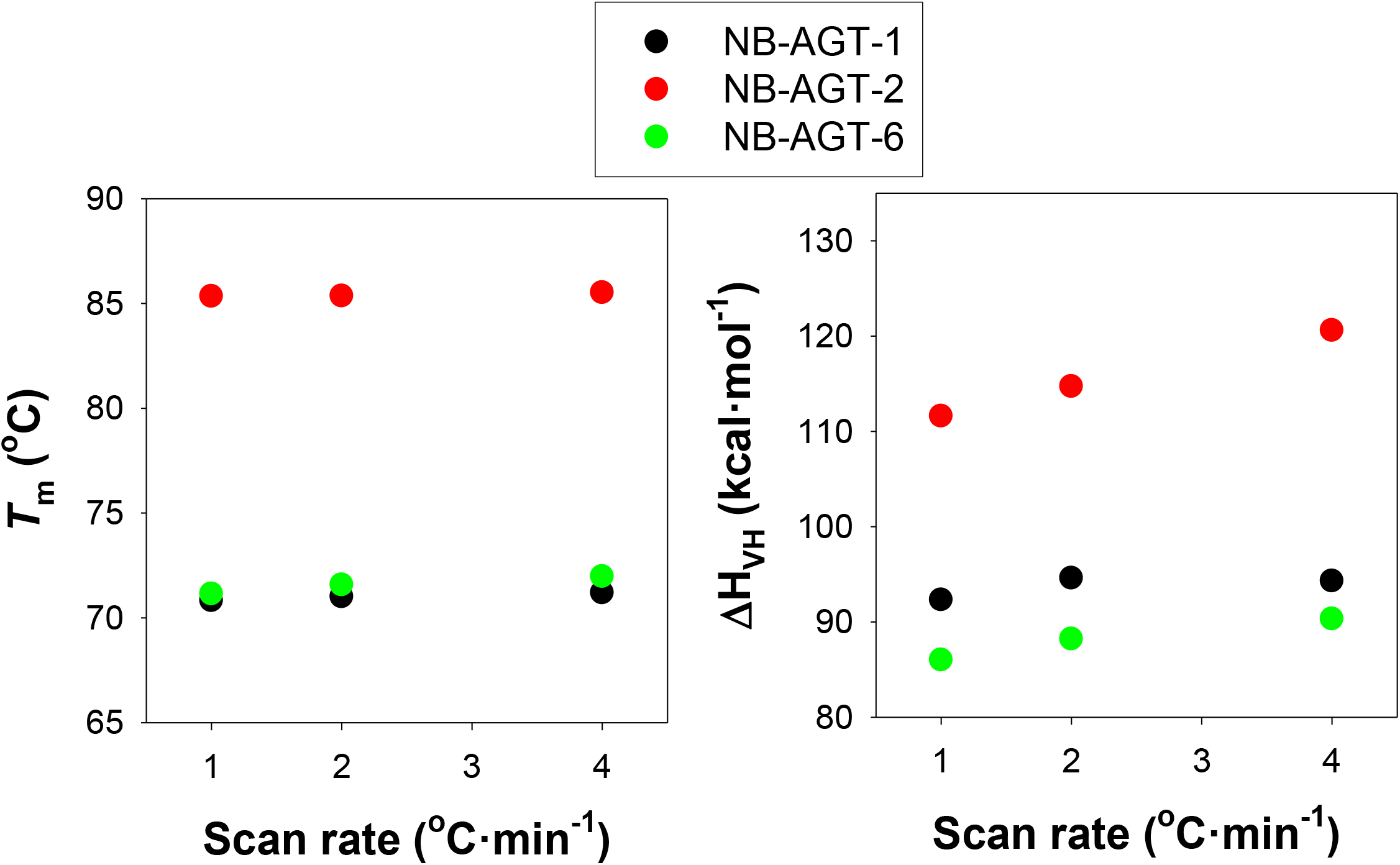
A two-state equilibrium thermal denaturation model by the weak scan-rate dependence of the thermal unfolding of NB-AGT-1, NB-AGT-2 and NB-AGT-6. Left panel shows the *T*_m_ for each NB-AGT determined using a two-state reversible unfolding model at different scan rates (Equation 1). Right panel shows the effects of scan rates on the Van’t Hoff enthalpies (which describe the dependence of unfolding equilibrium constant on temperature). The errors from fittings cannot be seen since these are smaller than the symbols. Data are from a single experiment at each scan rate.

### Isothermal denaturation by GdmHCl

GdmHCl denaturation of NB-AGT were carried out by mixing protein solutions with different concentrations of GdmHCl (typically in the 0-6 M range, the actual concentration determined using refractive index measurements). Experiments were carried out in 50 mM K-phosphate pH 7.4 using 5 µM of NB-AGTs. Samples were incubated at 4°C for 24 h and then these were stepwise incubated for 20 min at 18, 25, 32, 39 and 46°C. Once samples equilibrated, protein unfolding was monitored by fluorescence spectroscopy at that temperature, using an excitation wavelength of 295 nm and emission in the range 320-380 nm (both with 5 nm slits). Fluorescence measurements were carried out using 3×3 mm quartz cuvettes (Hellma Analytics, LineaLab, Badalona, Spain) in a Cary Eclipse spectrofluorimeter (Agilent Technologies, Madrid, Spain). Blanks in the absence of protein were routinely measured and subtracted. To monitor denaturation, we used two different spectral properties: the ratio of emission intensities at 365 and 335 nm (I_365_/I_335_) and the spectral center of mass (SCM, in nm, in the 320-380 nm range) the latter calculated using equation 3:

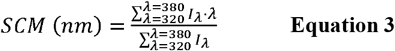

Where λ is the emission wavelength and I_λ_ is the emission intensity at that wavelength. To extract equilibrium denaturation parameters, we used a two-state unfolding equilibrium model [26](Equation 4):

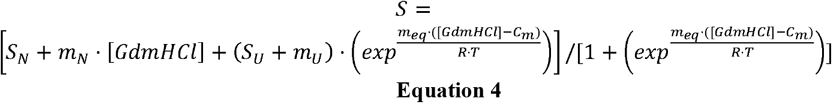

where *S* is the experimental spectral feature as a function of the [GdmHCl], *S*_N_ and *S*_U_ are the fitted spectral features for the native and unfolded state baselines at 0 M GdmHCl, respectively, *m*_N_ and *m*_U_ are the slopes of the native and unfolded state baselines, *m*_eq_ describes the unfolding cooperativity, *C*_m_ is the half-denaturation GdmHCl concentration, *R* is the ideal gas constant and *T* is the experimental temperature (in K). This model provides an excellent description of chemical denaturation of NB-AGT (Figure 4) as well as other NB [14]. To yield more accurate *m*_eq_ and *C*_m_ values, data for a given NB-AGT and temperature using both spectral features were simultaneously (globally) fitted. The product of *C*_*m*_ and *m*_eq_ provides the Δ*G* at a given temperature from GdmHCl induced unfolding profiles.

### Stability curves

To carry out simultaneous analyses of denaturation at low temperatures (from GdmHCl denaturation profiles) and high temperatures (in the absence of denaturation) we use an expression for the Gibbs-Helmholtz integrated equation for a two-state folder (Equation 5)[27]:

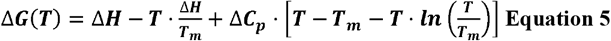

Where Δ*G*(*T*) is the temperature-dependent unfolding free Gibbs Energy, *T* is the experimental temperature, Δ*H*_cal_ is the calorimetric unfolding enthalpy change at the *T*_m_, *T*_m_ is the temperature at which the Δ*G*=0, Δ*C*_p_ is the change in heat capacity between the folded and unfolded states. Δ*G*(*T*) is calculated from DSC scans using a version of the Van’t Hoff equation (Equation 2). Good fits were obtained for NB-AGT-1 and 6 but not for NB-AGT-2.

### Structural models

To generate structural models of the three nanobodies we used Alpha-Fold 2 [28](https://colab.research.google.com/github/sokrypton/ColabFold/blob/main/AlphaFold2.ipynb), using the amino acid sequence of our NB-AGT and the structure with PDB code 4ZG1 as template. In this modeling we used default parameters and the higher ranked model for each NB-AGT. Accessible surface area in these models were calculated using GetArea (https://curie.utmb.edu/getarea.html) [29]. Multiple sequence alignment was carried out using Clustal Omega (https://www.ebi.ac.uk/jdispatcher/msa/clustalo).

## Results and discussion

### Global unfolding of NB-AGT by temperature follows a two-state equilibrium model

We first carried out thermal denaturation assays by DSC, and checked for their reversibility to focus on differences in thermodynamic stability related to changes in thermostability (rather than investigating aggregation propensities at very high, non-physiological and non-pharmaceutically related temperatures, that often required undesired long Temperature-extrapolations)[18,19]. As we show in Figure 1, the reversibility of the thermal transitions for NB-AGT 1, 2 and 6 is about 50%, consistent with previous studies showing that NB thermal unfolding is typically of ∼50-100% [14,24]. Additionally, the high symmetry of the peaks and the excellent fits to a simple two-state denaturation model, support that any source of irreversibility hardly affects the experimental scans and an appropriate analysis using equilibrium thermodynamics can be performed. Furthermore, the marginal effects of the scan rate on the thermodynamic parameters derived from the analysis using this reversible model (Figure 2) additionally support our DSC analysis. We considered an exception the determination of the unfolded heat capacity that would require heating up the samples to higher temperatures and would likely be affected more severely by irreversible processes [30]. Thus, we did not include the unfolding heat capacity change (Δ*C*_p_) in the model (see Equation 1). Instead, we estimated this parameter by performing DSC experiments in the presence of low concentrations of GdmHCl (Figure 3).

**Figure 3.**
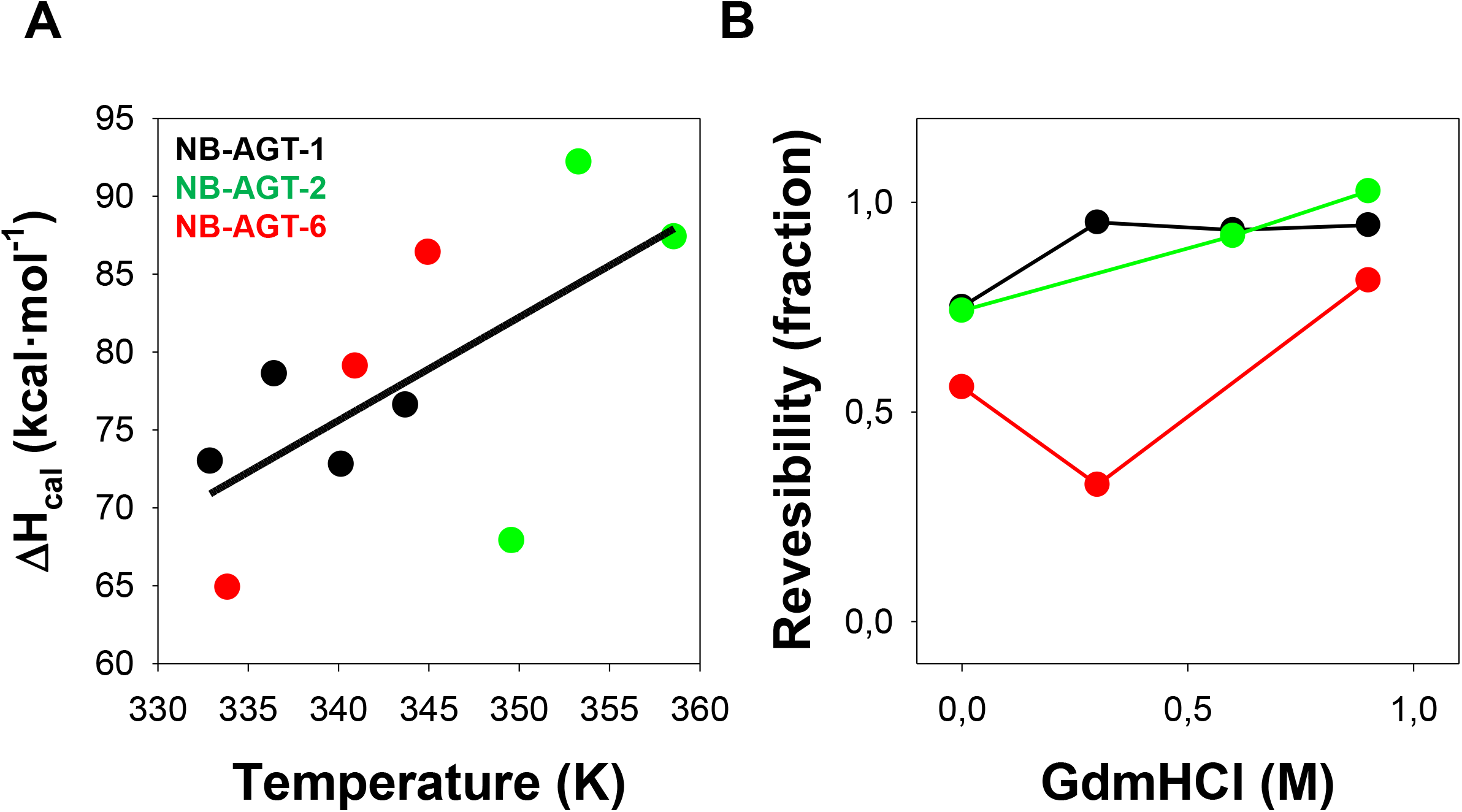
Thermal denaturation of NB-AGTs with low concentrations of GdmHCl. A) Calorimetric enthalpy (Δ*H*_cal_) as a function of the *T*_m_. The straight line is an overall fit that provides an unfolding change in heat capacity of 0.80±0.21 kcal·mol^-1^·K^-1^. The Δ*C*_p_ values predicted for proteins of this size are of ∼1.8 kcal·mol^-1^·K^-1^ [31]. B) Reversibility (as calculated from calorimetric enthalpies of upscans and rescans) of the thermal denaturation NB-AGT as function of denaturant concentration. Note that with 0.9 M of denaturant reversibility is almost complete. Symbols correspond to NB-AGT-1 (black), NB-AGT-2 (green) and NB-AGT-6 (red).

**Figure 4.**
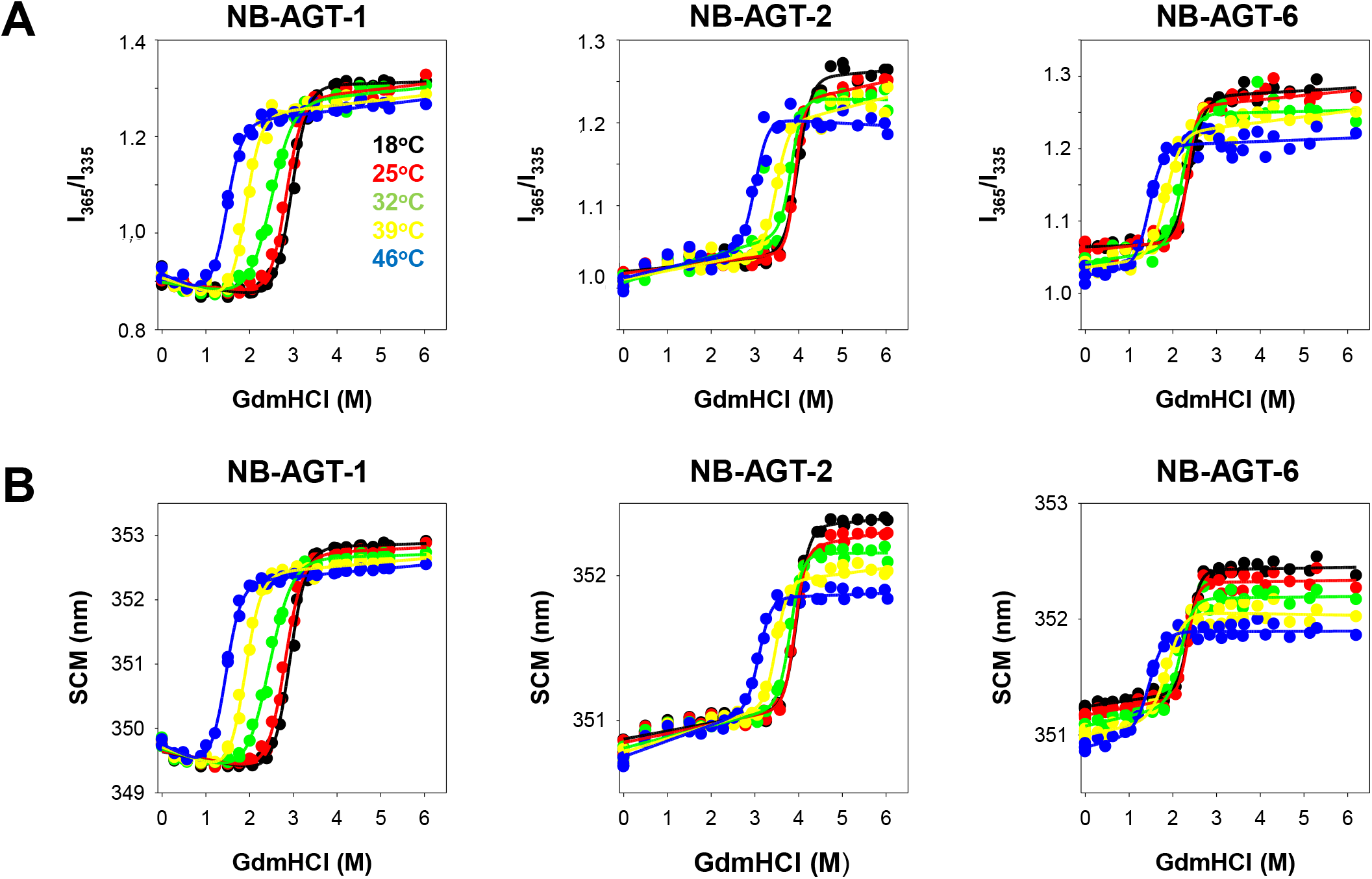
GdmHCl induced isothermal denaturation of NB-AGT at different temperatures. Different panels show the results from these experiments using different feature of the intrinsic emission fluorescence spectra (A, the I_365_/I_335_ ratio; B, the SCM) for the indicated NB-AGT. Best fits using equation 4 are shown as lines. The results for different temperatures are shown according to the color code.

It is evident that NB-AGT-2 is significantly more thermostable that NB-AGT-1 and 6 (about 15°C higher *T*_m_)(Figure 1 and Table 1). This higher thermostability could arise from either higher thermodynamic stability (at all temperatures), a higher temperature for maximal thermodynamic stability or from different temperature-dependencies of thermodynamic stability (e.g. different unfolding heat capacities, Δ*C*_p_)[27]. Importantly, reversibility tests as well as analysis of DSC transitions using a non-explicit two-state unfolding model (Table 1), and the lack of significant scan-rate dependence (Figure 2-3) supported that DSC analyses using a simple two-state unfolding model provide physically meaningful information on the unfolding free energy changes (Δ*G*) around the *T*_m_ (please, see the values for the Δ*H*_cal_/Δ*H*_VH_ very close to unity, consistent with a two-state unfolding behavior)(Table 1). Since the value of the Δ*C*_p_ for each NB-AGT cannot be extracted from these DSC analyses, we estimated an average of the Δ*C*_p_ = 0.8±0.2 kcal·mol^-1^ from the *T*_m_-dependence of Δ*H*_cal_ (Figure 3A; the *T*_m_ was gradually decreased for each NB-AGT by addition of low and non-denaturing GdmHCl concentrations).

**Table 1.** Equilibrium thermal denaturation parameters for the NB-AGT as determined by DSC. Parameters were determined using equations 1-2, that does not implicitly assume a two-state unfolding.

| NB-AGT | Equilibrium thermal unfolding parameters |  |  |  |  |
| --- | --- | --- | --- | --- | --- |
| | $T_m$ (°C) | $\Delta H_{cal}$ (kcal·mol<sup>-1</sup>) | $\Delta H_{VH}$ (kcal·mol<sup>-1</sup>) | $\Delta H_{cal}/\Delta H_{VH}$ | Reversibility |
| 1 | 70.65±0.01 | 83.3±0.2 | 92.3±0.3 | 0.90±0.01 | 47±1 % |
| 2 | 85.42±0.01 | 87.4±0.2 | 105.9±0.3 | 0.82±0.01 | 52±2 % |
| 6 | 71.68±0.01 | 77.1±0.2 | 94.1±0.3 | 0.82±0.01 | 50±1 % |

### Reversible chemical denaturation at different temperatures

In an attempt to determine the stability curve (Δ*G* vs T, equation 5) of NB-AGTs, we combined the DSC analyses (that monitored unfolding at high temperatures) with chemical denaturation analyses at lower temperatures (18-46°C). The use of GdmHCl as denaturant is justified because it causes reversible unfolding of NB and it is a stronger denaturant than urea (the later does not fully denature some NB)[14].

Chemical denaturation was followed by intrinsic tryptophan fluorescence, to use low protein concentrations and maximize reversible unfolding. The two spectral features (the ratio of intensities at 365 nm and 335 nm, I_365_/_335_, and the spectral center of mass, SCM) showed similar trends and a single transition, supporting that a two-state unfolding model is a reasonable description for the chemical denaturation of all NB-AGT investigated (Figure 4). Data suggested that the higher thermostability of NB-AGT-2 roots in a higher maximal Δ*G* at all temperatures temperatures tested (about 8-9 kcal·mol^-1^ at 25°C, Figure 5), whereas the temperature-dependence of the unfolding free energy (related to the unfolding heat capacity) seems to be quite similar. All *m*_eq_ were also similar, with average values (from different temperatures) of 3.1±0.4 (NB-AGT-1), 4.4±0.5 (NB-AGT-2) and 4.1±0.8 (NB-AGT-6) kcal·mol^-1^·M^-1^ supporting that their native and unfolded states are quite similar overall [31]. Actually, the theoretical *m*_eq_ value for proteins of these sizes is 3.3±0.1 kcal·mol^-1^·M^-1^ based on structure-energetics correlations [31]. When these average values of *m*_eq_ for a given variant were used, we observed that at their maximum, the Δ*G* value is ∼ 18 kcal·mol^-1^ for NB-AGT-2 and ∼ 9-10 kcal·mol^-1^ for NB-AGT-1 and 6 (Figure 5). Estimations of the Δ*C*_p_ could also be determined from stability curves (Figure 5) with values of 1.1-1.3 kcal·mol^-1^·K^-1^ (Figure 5). These values were somewhat higher that those obtained from DSC experiments in the presence of low GdmHCl concentrations (0.8±0.2 kcal·mol^-1^·K^-1^; Figure 3A). It is also worth noting that reversibility of thermal denaturation of all NB-AGT were enhanced with low concentrations of GdmHCl (up to 80-100% reversibility; Figure 3B).

**Figure 5.**
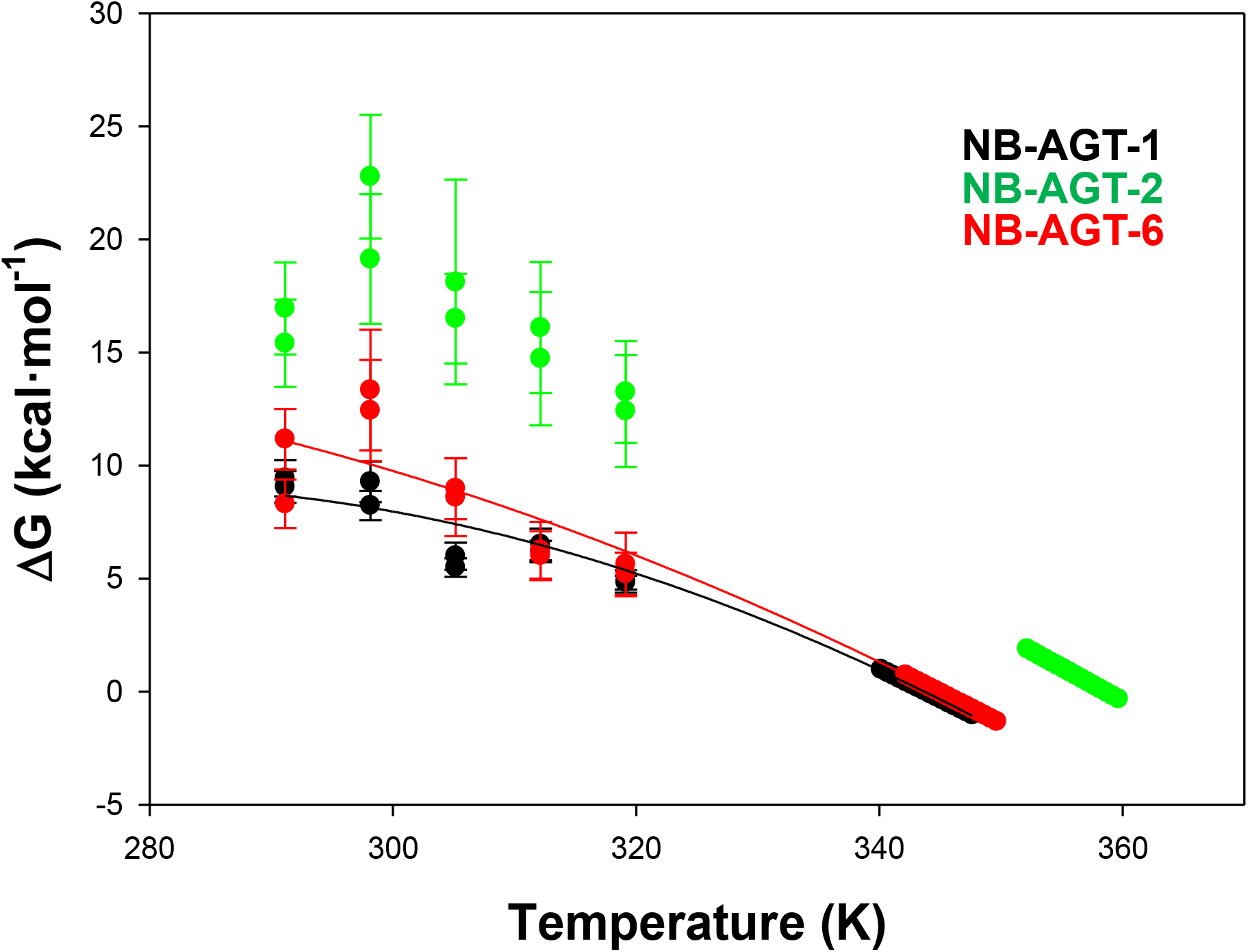
Stability curves built using chemical and thermal denaturation data. Data for NB-AGT-1 and 6 were successfully fitted to equation 5 providing Δ*C*_p_ values of 1.30±0.05 and 1.07±0.04 kcal·mol^-1^·K^-1^. Data for NB-AGT-2 did not provide a good fit. For fittings, we used the *T*_m_ and Δ*H*_cal_ for each variant as fixed parameters (Table 1).

### Similar predicted structures for the NB-AGT

To get an estimation of the structural differences between the three NB-AGT, we have used the recently developed algorithm Alpha-Fold 2 and the sequences of the NB-AGT. A structural alignment of the three NB-AGT models with the highest rank showed a very similar overall structure (pairwise Cα RMSD of 0.25 Å for AGT-1 vs. AGT-2 and 0.39 Å) (Figure 6A). Visual inspection of this superposition showed a clear difference in the HVL-3 (belonging to the CDR-3)(Figure 6A). A multiple sequence alignment showed high conservation (identity of 60.6-64.0%, including similar residues of 70.5%-74.4%), particularly in the β-sheets, whereas the VHL1-2 and surroundings (i.e. CDR1-2) showed a moderate degree of conservation and the VHL-3 (and CDR3) displayed virtually no conservation (Figure 6B). Calculation of ASA in the modeled structure showed very similar solvent exposure of different structural regions, with high exposure of the three HVL (Figure 6C).

**Figure 6.**
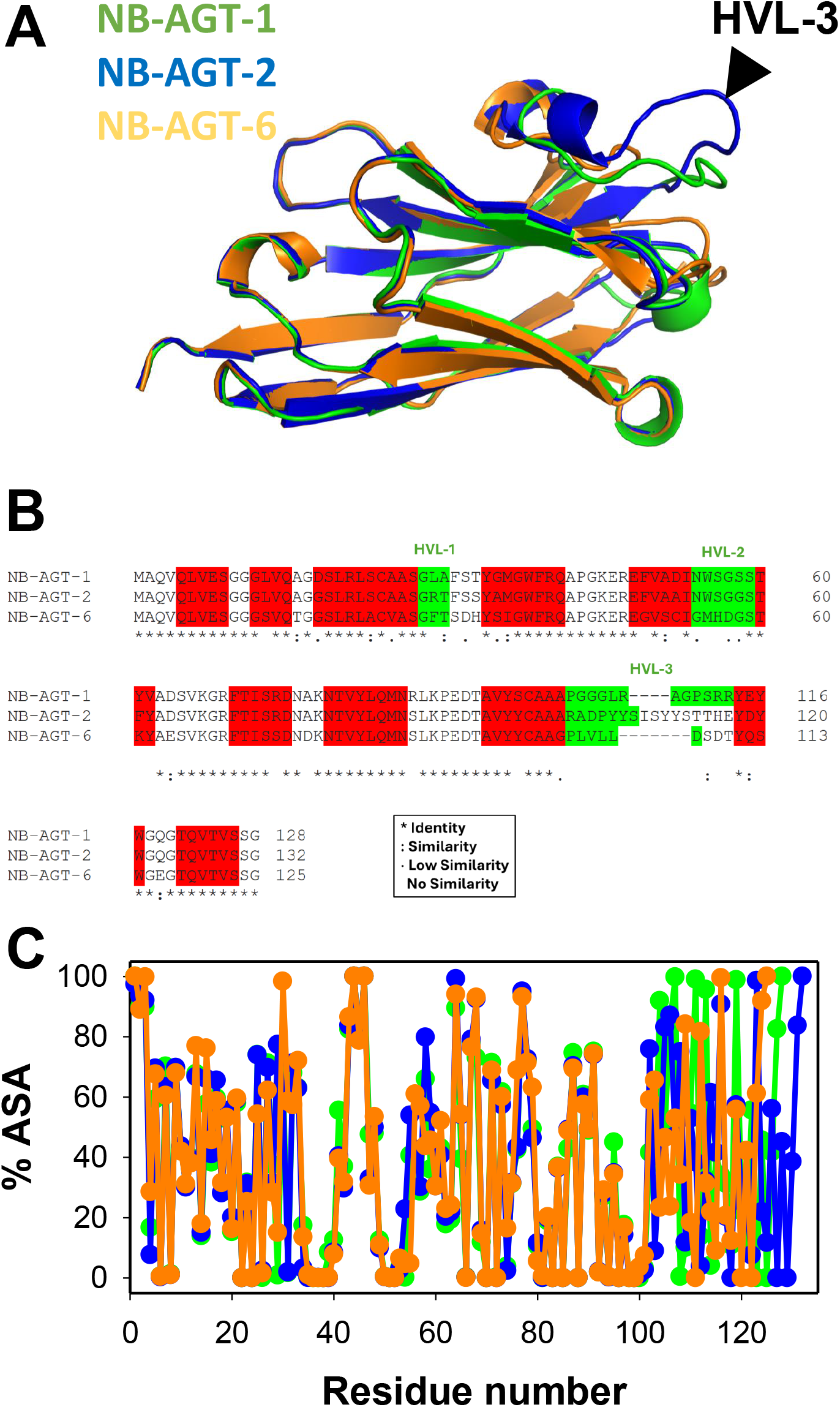
Structural models and sequence alignment for NB-AGTs. A) Highest-scored structural models generated by Alpha-Fold 2 (Mirdita et al., 2022) for NB-AGT-1 (in green), NB-AGT-2 (in blue) and NB-AGT-6 (in orange). B) Sequence identity for a multiple alignment (using ClustalOmega; https://www.ebi.ac.uk/Tools/msa/clustalo/) of NB-AGT-1, -2 and -6. The identity or similarity between amino acids is shown using the symbols found in the inset. Amino acid belonging to a β-sheet structure are highlighted in red, and those belonging to HVLs are highlighted in green, all these based on the structural alignment shown in panel A. C) The accessible surface area (% ASA; average of the three best-scored models generated by Alpha-Fold 2 for each NB-AGT) determined using GetArea; https://curie.utmb.edu/getarea.html) for NB-AGT. The color code in panel C is the same that in panel A. Note that in all three panels, the highest divergence is found in the HVL-3.

In conclusion, our study shows that reversible chemical unfolding at storage or physiological temperatures can be described by equilibrium thermodynamics and correlate with thermal stability for our NB-AGT, supporting the results of previous studies with other NB [14,24]. It would be interesting to carry out kinetic denaturation experiments using GdmHCl to know whether unfolding (as a *k*_u_) and aggregation (as *k*_agg_) rate constants also correlate with thermodynamic stability at storage or physiological temperatures. We must note that these two rate constants may differ by order of magnitudes at the same temperature [15,32]. Therefore, we must be cautious when data obtained at very harsh conditions (close to the apparent *T*_m_) are used to compare the thermal stability with the thermodynamic/kinetic stability at lower (storage or physiological) temperatures [18,19,33] for a given NB.

## Acknowledgements

This work was supported by Consejería de Economía, Conocimiento, Empresas y Universidad, Junta de Andalucía [Grant number P18-RT-2413].

## Author contributions

A.L.P was responsible for conceptualization. A.G-M purified proteins and carried out experiments. A.L.P. carried out data analysis. E.S. provided reagents. A.L.P. wrote the paper and all authors contributed to editing and revising the manuscript.

## Abbreviations

AGT-LM.: minor allele of the human alanine glyoxylate aminotransferase
BB.: binding buffer
CDR.: complementarity-determining region
*C*_m_.: concentration of GdmHCl for half-unfolding
*C*_p,app_.: unfolding apparent heat capacity
DSC.: differential scanning calorimetry
GdmHCl.: guanidinium hydrochloride
IPTG.: isopropyl β-D-1-thiogalactopyranoside)
HEPES.: N-(2-Hydroxyethyl)piperazine-N’-(2-ethanesulfonic acid)
HPLC/ESI-MS.: high performance liquid chromatography coupled to electrospray ionization mass spectrometry
HVL.: hypervariable loop
I.: fluorescence emission intensity
IMAC.: immobilized metal affinity chromatography
*K*.: unfolding equilibrium constant
ku.: unfolding rate constant
*k*_agg_.: aggregation rate constant
LB.: Luria-Bertani
LBK.: LB with kanamycin
*m*_eq_.: chemical unfolding cooperativity
NB.: nanobody
NB-AGT-1, 2 and 6.: NB 1, 2 and 6 raised against AGT-LM
SDS-PAGE.: Polyacrylamide gel electrophoresis in the presence of sodium dodecyl sulphate
*T*_m_.: temperature at which *K*=1
Δ*C*_p_.: unfolding heat capacity change
Δ*G*.: unfolding free energy change
Δ*H*_cal_.: unfolding calorimetric change in enthalpy
Δ*H*_VH_.: unfolding Van’t Hoff enthalpy.

